# Comparative metaproteomics demonstrates different views on the complex granular sludge microbiome

**DOI:** 10.1101/2022.03.07.483319

**Authors:** Hugo B.C. Kleikamp, Dennis Grouzdev, Pim Schaasberg, Ramon van Valderen, Ramon van der Zwaan, Roel van de Wijgaart, Yuemei Lin, Ben Abbas, Mario Pronk, Mark C.M. van Loosdrecht, Martin Pabst

## Abstract

The tremendous progress in sequencing technologies has made 16S amplicon and whole metagenome sequencing routine in microbiome studies. Furthermore, advances in mass spectrometric techniques has expanded conventional proteomics into the field of microbial ecology. Commonly referred to as metaproteomics, this approach measures the gene products (i.e., proteins) to subsequently identify the actively-expressed metabolic pathways and the protein-biomass composition of complete microbial communities.

However, more systematic studies on metaproteomic and genomic approaches are urgently needed, to determine the orthogonal character of these approaches. Here we describe a deep, comparative metaproteomic study on the complex aerobic granular sludge microbiome obtained from different wastewater treatment plants. Thereby, we demonstrate the different views that can be obtained on the central nutrient-removing organisms depending on the ‘omic’ approach and reference sequence databases. Furthermore, we demonstrate a ‘homogenized’ Genome Taxonomy Database (GTDB) that subsequently enables a more accurate interpretation of data from different omics approaches. Ultimately, our systematic study underscores the importance of metaproteomics in the characterization of complex microbiomes; and the necessity of accurate reference sequence databases to improve the comparison between approaches and accuracy in scientific reporting.

## INTRODUCTION

Microbial communities play a central role in the global biogeochemical cycles and their close association with humans has a direct impact on health and disease [1–5]. Moreover, microbial communities are increasingly used in biotechnology and engineering such as for the degradation and removal of pollutants from wastewater and soils, or for the production of novel materials, greener chemicals, or energy to support a bio-based society [6–11]. Of more recent interest are also synthetic and engineered communities with the aim to enable fundamentally novel applications [12]. The complex nature of microbial interactions, however, still hampers the design of specific functions in such environments [12, 13]. Nevertheless, the need to better understand global microbial processes and the desire to harness microbial communities for industrial applications asks for methods that resolve the taxonomic composition and their underlying metabolic pathways. Therefore, systems biology approaches that provide molecular level information from such complex environments become increasingly important across biotechnology and microbial ecology.

The emergence of next-generation sequencing (NGS) technologies has enabled large-scale genomic studies of microbial communities directly from their natural environments. The simplest of these approaches is 16S rRNA gene sequencing (commonly referred to 16S amplicon sequencing). 16S rRNA genes are highly-conserved between different bacteria and archaea and, thus, are widely-used in taxonomic profiling of environmental communities [14–19]. However, this approach suffers from the variable 16S gene copy numbers [20–22] and primer efficiencies across microbes [23, 24]. Furthermore, metabolic functions are only inferred from prior taxonomic knowledge and thus remain purely predictive [25, 26]. Alternatively, whole metagenome (‘shotgun’) sequencing (often referred to as metagenomics) targets the complete genomes of all community members. This approach provides a high taxonomic resolution and it discloses the metabolic potential of individual community members [27–29]. The obtained metagenome, however, may not only encompass the active microbial population, but can also cover free DNA as well as DNA from dead and dormant microbes [30].

Advances in high-resolution mass spectrometry and the increased ease to construct proteome sequence databases ultimately enabled deep proteomic studies of complete microbial communities. Consequently, metaproteomics has emerged as one of the most promising post-genomic approaches [31–38]. Most importantly, because metaproteomics measures the gene products (*i.e.,* proteins) it provides an orthogonal view on the microbial community. The obtained microbial composition correlates to the amount of protein (proteinaceous biomass) rather than to the number of cells, as obtained by metagenomics [35, 39]. Therefore, metaproteomic data resemble more closely the metabolic capacity of individual community members [40–42]. Moreover, metaproteomics allows to measure molecular level information such as protein modifications that cannot be obtained from genomic information alone [43, 44]. However, in contrast to DNA, proteins cannot be amplified prior to analysis, and peptide sequencing is performed consecutively (or only at low multiplexing level) rather than (all sequences) in parallel. Therefore, the depth of information that can be obtained by metaproteomics is dependent on the taxonomic complexity and the mass spectrometric effort taken to sequence the sample [31, 32, 45]. The dependency of metaproteomic performance on community complexity has been investigated more in detail by Lohmann and co-workers only recently [46].

From the many applications in industrial biotechnology, agriculture or medicine, microbial water treatment is perhaps one of the fastest growing areas. For example, the widely-used activated sludge wastewater treatment technology is a biological process that with the aid of a complex microbiome aims to purify wastewater [47, 48]. A recent advancement, known as aerobic granular sludge (AGS) technology, has the advantage of operating with reduced space and energy requirements [6, 27–30]. The microbes form dense granules following the production of extracellular polymeric substances [49–51]. Consequently, the granules allow a settling speed and a higher biomass density. In microbial wastewater treatment, several synergistic roles for nutrient removal have been identified that include phosphate-accumulating organisms (PAO), glycogen-accumulating organisms (GAO), nitrate-oxidizing bacteria (NOB), ammonia-oxidizing bacteria (AOB), and nitrate reducers (NR) [52–54]. Although microbial wastewater treatment has a long history, the exact molecular-level processes and the organisms that are involved in nutrient removal are still poorly understood [19, 55]. Therefore, determining the taxonomic composition of the core microbiome and the expressed metabolic functions are important in optimizing purification processes and developing better purification strategies.

Large-scale genome sequencing efforts on activated sludge established the wastewater microbiome specific database called ‘MiDAS’ (Microbial Database for Activated Sludge). The consortium uniformly applies full-length 16S rRNA gene sequencing to create a worldwide map of microbes present in activated sludge systems, with the aim to link organisms to nutrient-removal functions [38–40]. Information obtained from DNA and rRNA-based approaches, however, have been often found to contradict staining experiments or measured metabolic conversions [16, 56, 57]. This underlines the importance of employing additional (orthogonal) approaches – such as metaproteomics – when characterizing complex communities. Nevertheless, the lack of standardization within the ‘omics’ field makes a comparison of different experiments highly challenging. This was observed even for studies that were performed with the same types of omics approaches [58, 59].

For example, metagenomics experiments are commonly employed to construct protein sequence databases for metaproteomics studies in order to enable a deep sequence coverage and a high taxonomic and functional resolution. A comprehensive taxonomic classification of the metagenomic data, however, relies on accurate and complete reference sequence databases. Consequently, a potential large source of variation and inaccuracy derives already from the reference databases that have been used to taxonomically classify the metagenomic sequences. Different reference databases substantially vary in taxonomic coverage, sequence content and the nomenclature and employed phylogenies. Modern phylogenetic placement tools employ a range of methods such as 16S similarity, average amino acid identity and average nucleotide identity [60, 61]. The NCBI taxonomy, that is used for RefSeq and UniProtKB employs a mixture of historical taxonomies and modern placement methods and lacks a rank normalization that ultimately results in lineages with gaps in taxonomic annotations (further referred to as ‘gapped lineages’) [62, 63]. In addition NCBI taxonomies contain taxa that cluster groups of uncultured organisms (further referred to as ‘dump taxa’) [64]. Thus, the NCBI taxonomy is often not consistent with respect to true evolutionary relationships. Many taxa circumscribe polyphyletic groupings and there is an uneven application of ranks across the phylogenetic tree [65–67].

Standardized reference sequence databases with accurate taxonomies are therefore of utmost importance to accurately describe microbial diversity, enable data comparison between experiments and approaches, and communicate scientific data [65, 68]. The recently-established genome taxonomy database (GTDB) addresses these issues by using a set of conserved proteins and employing a placement method that normalizes taxonomic ranks based on relative evolutionary divergence [65, 69–71]. The GTDB taxonomy offers an objective, phylogenetically-consistent classification of prokaryotic species, and therefore enables a more accurate description of the taxonomic and metabolic diversity of a microbial community [65].

The Genome Taxonomy Database Toolkit (GTDB-Tk) supports the classification of draft bacterial and archaeal genomes [70]. However, this tool was developed for genome assemblies or metagenome-assembled genomes that are constructed by clustering related contigs into bins [72, 73]. The binning procedure, however, leaves for complex metagenomes usually a substantial fraction of unbinned sequences [59]. Consequently, assembled genomes often provide a substantially less complete sequence reference database compared to the alternative reads- or contig-based databases [74–78], which is a major drawback for metaproteomic studies. For that reason, database construction and taxonomic classification has been frequently performed on contigs or scaffolds, e.g. as demonstrated by the contig annotation tool (CAT) only recently [79].

In this study, we describe a deep metaproteomic characterization of the aerobic granular sludge microbiome obtained from different wastewater treatment plants, and systematically compare the observed taxonomic and metabolic profiles to the orthogonal DNA and rRNA-based approaches. Moreover, we demonstrate the application of a ‘homogenized’ genome taxonomy database and showcase the impact of divergent reference database content on the outcomes. Ultimately, this comparative omics study underscores the importance of orthogonal metaproteomics experiments when characterizing complex microbiomes.

## MATERIALS AND METHODS

### Sampling of aerobic granular sludge

Aerobic granular sludge (AGS) was collected from three different full-scale AGS wastewater treatment plants in the Netherlands: Dinxperlo (DX, plant 1), Garmerwolde (GW, plant 2) and Simpelveld (SP, plant 3). Each plant performed stable operation with simultaneous denitrification and phosphorous removal. AGS granules were sieved to select a size fraction with a diameter of approximately 2.0 mm. Granules were stored at −80”C until further processed.

### Protein extraction and proteolytic digestion

The collected granules were freeze-dried and ground with a mortar and pestle. Two hundred milligrams of acid washed glass beads (150–212 μm) and 350 μL of both TEAB and B-PER buffer were added to approximately 5 mg starting material. Bead beating was performed for 20 s (×3) with a 30 s pause between cycles. Samples were centrifuged and freeze/thaw cycles (×3) were performed by freezing the sample at −80°C and subsequently thawing at 95°C in a water bath. The samples were centrifuged, and the supernatant was collected. Protein precipitation was performed by adding TCA at a ratio of TCA to supernatant of 1:4. The samples were incubated at 4°C for 10 min. and then centrifuged at 14,000 r.p.m. for 5 min. The pellets were washed with 200 μL ice-cold acetone. The protein pellets were reconstituted in 250 μL 6 M urea and the protein extracts were then reduced with 10 mM dithiothreitol (DTT) for 60 min. at 37°C. Next, the samples were alkylated with 20 mM iodoacetamide (IAA) and incubated in the dark at room temperature for 30 min. Thereafter, the samples were diluted with 200 mM ammonium bicarbonate (AmBiC) to <1 M urea. Finally, sequencing-grade trypsin was added (Promega) at an approximate enzyme to protein ratio of 1:50 and incubated at 37°C overnight. The obtained peptides were purified by solid-phase extraction using Oasis HLB solid-phase extraction well plates (Waters) according to the protocol provided by the manufacturer. Purified peptide fractions were then dried in a SpeedVac concentrator, reconstituted in aqueous 0.1% TFA and separated (according to the instructions supplied by the manufacturer) into 8 fractions using the Pierce high pH reversed-phase fractionation kit (Thermo Scientific). For plants 2 (DX) and 3 (GW) the fractions 2+6, 3+7, 4+8 were combined. The obtained samples were dried in a SpeedVac concentrator and dissolved in water containing 3% acetonitrile and 0.1% formic acid, resulting in 8 fractions for plant 1 (DX), 4 fractions for plant 2 (GW) and 3 (SP). The approximate concentration of the protein digest was determined using a NanoDrop micro-volume spectrophotometer.

### Shotgun metaproteomic analysis

Briefly, the prepared fractions were analyzed by injecting approx. 300 ng proteolytic digest using a one-dimensional shotgun proteomic approach on a nano-liquid-chromatography system consisting of an EASY nano-LC 1200 equipped with an Acclaim PepMap RSLC RP C18 separation column (50 μm × 150 mm, 2 μm and 100 Å) coupled to a QE Plus Orbitrap mass spectrometer (Thermo Scientific, Germany). The flow rate was maintained at 350 nL/min using as solvent A water containing 0.1% formic acid, and as solvent B 80% acetonitrile in water and 0.1% formic acid. The Orbitrap was operated in data-dependent acquisition mode acquiring peptide signals at 70 K resolution and a max IT of 100 ms, where the top 10 precursor ions were isolated by a 2.0 *m/z* window with an 0.1 *m/z* isolation offset, and fragmented at an NCE of 28. The AGC target was set to 2e5 at a max. IT of 75 ms and 17.5 K resolution. Mass peaks with unassigned charge state, singly, 7 and >7, were excluded from fragmentation. For the prepared fractions from plants 2 (GW) and 3 (SP) analysis in duplicates was performed using a linear gradient from 5% to 28% solvent B for 115 min and finally to 55% B for additional 60 min. The individual fractions obtained from plant 1 were analysed by single injections using a short linear gradient from 6% to 26% solvent B for 45 min and finally to 50% B over additional 10 min.

### Processing of metaproteomic raw data

Mass spectrometric raw data (obtained from the fractions of each plant) were combined and analysed using PEAKS StudioX by either database searching against the metagenomic-constructed databases from predicted ORFs, or by *de novo* sequencing as quality control and to estimate the percentage of eukaryotic sequences [39, 80]. Redundant sequences in the constructed databases were removed by employing a local installation of CD-HIT [81]. Database search was performed by including cRAP protein sequences (https://www.thegpm.org/crap/), setting carbamidomethylation (C) as fixed and oxidation (M) and deamidation (N/Q) as variable modifications, allowing up to 2 missed cleavages and 2 variable modifications per peptide. Peptide-spectrum matches were filtered against 1% false discovery rate and protein identifications with ≥2 unique peptides were considered as significant. Taxonomic annotation of database-matched peptide sequences was achieved by determining the lowest common ancestor (LCA) using the taxonomic classification obtained for the contigs (see below for taxonomic classification of metagenomics data). Metabolic annotation with KEGG orthologies was performed using BlastKOALA [82]. Moreover, WEBMGA [83] was used to annotate Clusters of Orthologous Groups (COGs) and protein families (PFAMs and the complementary TIGRFAM terms). DIAMOND v2.11 [84] was used to annotate ORFs with UniprotKB genes.

### DNA extraction and sequencing

Extraction of DNA for both 16S rRNA gene sequencing and shotgun metagenomics was performed using a DNeasy UltraClean Microbial Kit (Qiagen, Germany), and the extracted DNA was quantitated with a Qubit fluorometer. 16S rRNA gene amplification was performed by Novogene (Novogene Co., Ltd., China) by amplifying V3–V4 regions with 341F, 806R primers. Sequencing of 16S rRNA genes and shotgun metagenomics was achieved with paired-end reads on an Illumina NovaSeq platform.

### Processing of 16S rRNA raw sequencing data

Standard read preparation including demultiplexing, trimming and assembly was performed by Novogene (Novogene Co., Ltd., China). Cleaned reads were used to select amplicon sequence variants (ASVs) with Usearchv11 command -unoise3 [85]. To improve ASV selection, the data set was padded with additional sample sets containing different granule size fractions from each water treatment plant: flocs, >0.2, >0.7, >1.0 mm (data not shown). Taxonomic annotation was performed using QIIME2 [86] with trained V3–V4 classifiers. As a comparison, 16S rRNA sequences were annotated with GTDB representative of small subunit ribosomal RNA (ssu rRNA) sequences, Midas 3.7 flASVs, and a SILVA NR99 v138 pre-trained V3–V4 classifier [87]. To compare the effects of database homogenization, GTDB r202 complete 16S (ssu all) and representative (ssu reps) were analyzed with all sequences and full-length sequences of >1200 base pairs.

### Processing of metagenomic raw sequencing data

Reads were assembled for all samples using metaSPAdes v3.14.0 at default settings [88]. Prodigal v2.6.3 was employed as a gene caller to identify open reading frames (ORFs) [89]. DIAMOND v2.11 was used to align ORFs with parameters -fast -top 10 -e 0.001 (with otherwise default parameters) to protein databases of GTDB r202, from Uniprot release 2021 03: UniProtKB, Swiss-Prot UniRef100, UniRef90, UniRef50, and from NCBI RefSeq protein and RefSeq protein non-redundant release 205 [84]. The contig-level taxonomic classification was furthermore performed based on the ‘CAT’ approach published by Meijenfeldt et al., 2019 [79]. Firstly, the taxonomy of each ORF was determined by lowest ancestor analysis of the top Diamond hits followed by constructing a consensus lineage for each contig from the annotated ORFs. Adjustments compared to the original CAT approach were performed with the objective to maximize genus level annotations. Details about the adjusted approach is can be found in the supplementary information material and Figures S1– 2. Preprocessing of sequences obtained from the genome taxonomy database (GTDB) for the use with Diamond as well as the ‘protein LCA’ of the diamond alignments were performed with the python tools available via https://github.com/hbckleikamp/GTDB2DIAMOND. Reformatting of the sequences for the use with QIIME were performed with the python tools available via: https://github.com/hbckleikamp/GTDB2QIIME. The metagenome coverages were as estimated by Bowtie 2 v2.3.5.1 and QualiMap 2 v2.2.2 [90, 91] where the reads were first mapped to individual scaffolds using Bowtie and the obtained BAM file were analysed using QualiMap. The average depth of sequencing coverage was determined according to LN/G (L= length or read, N = number of reads and G = genome length) [92], which values were furthermore summed for every taxonomic identifier for compositional analysis as shown in the bar graphs.

### Homogenization of GTDB

The ‘homogenized’ GTDB protein reference sequence database was constructed from organisms that are also represented in the ‘GTDB ssu reps’ (small-subunit ribosomal RNA database) and that contained ‘full length’ 16S rRNA sequences (the cutoff to define ‘full length’ sequences was set to >1200 base pairs).

### Comparative analysis of taxonomic classifications and annotation coverage obtained by employing different databases

The annotation differences obtained from using the different reference sequence databases was visualized with Sankey flow diagrams. The nomenclature differences between GTDB and the other databases such as NCBI, UniprotKB, SILVA or MiDAS however challenges a comprehensive comparison. Therefore, taxonomic nomenclatures were ‘unified’ using auxiliary conversion tables obtained from https://data.gtdb.ecogenomic.org/releases/latest/. If taxonomic names matched at least 3 out of 4 times, the name was changed to the respective name reported in GTDB. Furthermore, ‘Candidatus’ prefixes and GTDB unique suffixes such as ‘Firmicutes_A’, ‘Firmicutes_B’, were removed. Gaps in the taxonomic lineage annotations were ‘bridged’. The representation of the taxonomic abundance in the graphs is based on total ASV counts for 16S amplicon sequencing data, (summed) depth of sequencing coverage of related contigs for metagenomics data, and total number of peptide-to-spectrum matches (PSMs) for metaproteomics data. The employed conversion tables (NCBI and SILVA to GTDB, and vice versa) can be found in the supplementary information (SI-Excel-1–3).

### Shared biomass fraction, diversity, richness and evenness

The shared biomass (or genera) was determined between taxonomies that were observed by at least two techniques (=non-unique taxa), and which taxa further were present at >3% abundance (compared to total abundance of the non-unique taxa within one technique). Taxa which express central nutrient-removing genes were included into the evaluation regardless their abundance. Diversity, richness, evenness and shared biomass were determined after uniformly applying an abundance cut-off of 0.1% in order to homogenize data treatment across the three techniques. Richness corresponds to the number of unique taxa. Simpson’s evenness and Shannon’s diversity were calculated using the Python functions ‘skbio.diversity.alpha.simpson_e(X)’ and ‘skbio.diversity.alpha_diversity(‘shannon’,X)’ which are part of the skbio Python package (http://scikit-bio.org). Determination of the abundance of taxa was based on total ASV counts for 16S amplicon, summed depth of contigs in metagenomics, and the total number of peptide-to-spectrum matches for metaproteomics.

### Functional classification, between technique abundance differences and COG term enrichment analysis

The total abundance was renormalized to a subset of non-unique taxa that showed an abundance of >3%, or that contained nutrient-removal genes. The between technique absolute abundance difference (*x-y*) and percent abundance difference (*x* – *y*)/(((*x* + *y*)) / 2) was then determined for every genus. The functional analysis and classification was performed by integrating KEGG, COG, PFAM, TIGRFAM and UniprotKB genes (for NXR). Two manually-curated sub-classifications were added to the COG system; ‘nitrogen metabolism’ (based on KEGG pathways) and ‘porin’ that includes beta-barrel proteins. Between method COG term enrichment was determined by comparing PSMs from metaproteomic experiments to read counts (‘summed sequencing depth’) from metagenomics experiments.

### Raw data availability

The mass spectrometry proteomics raw data have been deposited in the ProteomeXchange consortium database with the dataset identifier PXD030677. Raw sequencing data have been made available through the NCBI Sequence Read Archive (SRA) under accession number: SRP352708. The BioProject accession number is PRJNA792132.

## RESULTS

### A deep comparative metaproteomic study on the core microbiome of granular sludge

Here, we demonstrate a deep comparative metaproteomic study by simultaneously applying metaproteomics, metagenomics and 16S rRNA sequencing to the same granular sludge microbiome (with uniform 2 mm granule size). In addition to the different approaches, we systematically investigated the impact of employing different reference sequence databases on the obtained taxonomic profiles and metabolic functions (Figure 1). A uniform database with an accurate taxonomy is of greatest importance in order to improve comparability between different ‘omics’ approaches and to accurately capture the microbial diversity [65, 68]. The genome taxonomy database (GTDB) uses a set of conserved proteins to normalize taxonomic ranks based on relative evolutionary divergence with the aim to provide an objective, phylogenetically consistent classification of prokaryotes [65, 69–71]. The Genome Taxonomy Database Toolkit (GTDB-Tk) furthermore enables to efficiently classify bacterial and archaeal draft genome assemblies [70, 72, 73]. However, in metagenomics, the clustering and binning of contigs into individual genomes commonly results in a substantial number of unbinned fractions [77, 78]. This can significantly bias the taxonomic representation towards the more abundant organisms in a community.

**Figure 1:**
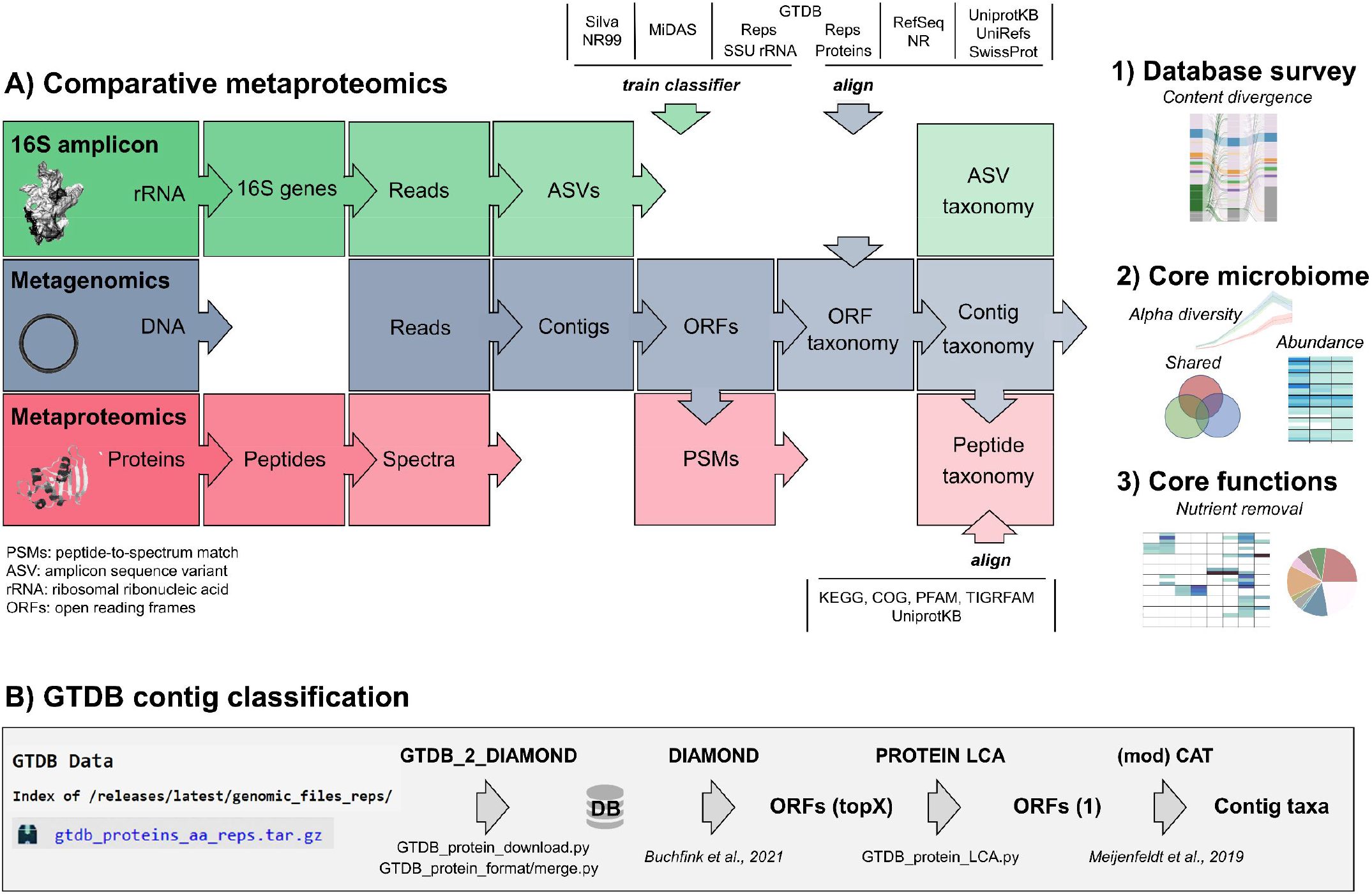
**A)** The graph shows a workflow diagram of ‘omics’ techniques and reference sequence databases used to perform a deep, comparative metaproteomics study on the complex microbiome of aerobic granular sludge. In addition to metaproteomics (red), the same microbiomes were analysed via orthogonal metagenomics (blue), and 16S rRNA amplicon sequencing (green). For 16S rRNA gene sequencing the V3–V4 region was amplified. Furthermore, the amplicon sequencing variants (ASVs) were determined and compared to small subunit ribosomal RNA (SSU rRNA) sequence databases for taxonomic classification (ASV taxonomy). For whole metagenome sequencing (metagenomics) reads were assembled into contigs. Identified ORFs were furthermore aligned to reference sequence databases for taxonomic classification (ORF taxonomy) to ultimately provide a taxonomic classification for the contigs (contig taxonomy). Metaproteomics was performed by analyzing the tryptic peptides using a shotgun proteomics approach. Peptide spectra were subsequently analysed using the protein sequences (ORFs) identified from the metagenomics experiments. Taxonomic classification was aligned to the taxonomies determined for the respective contigs. In addition to the different omics approaches, a range of different reference sequence databases were used for taxonomic classification. The obtained outcomes were finally evaluated for the 1) impact of reference sequence content divergence on obtained taxonomic profiles, 2) core microbiome that was consistently observed by all approaches, as well as the shared microbiome between the different treatment plants, and finally for the 3) expressed key functions as determined by metaproteomics. **B)** The scheme outlines the taxonomic classification of contigs using different reference sequence databases, as exemplified for the genome taxonomy database (GTDB). Reference sequences (protein reps) were downloaded from GTDB (https://gtdb.ecogenomic.org/downloads/), merged into a single file and reformatted to allow the use with the sequence aligner Diamond [84]. A consensus lineage was furthermore determined using a lowest common ancestor (LCA) algorithm, and finally, a contig-level lineage (taxonomy) was determined using a modified version of the CAT tool algorithm (details concerning the modified CAT algorithm are detailed in the SI-doc, chapter 1) [79]. The taxonomies assigned to individual contigs were subsequently used to classify the peptide-to-spectrum matches (PSMs) obtained by metaproteomics. The established python codes are openly accessible via Github (see methods section). To support comparison between the different approaches, a homogenized Genome Taxonomy Database (GTDB) was constructed from organisms represented in GTDB ssu reps and from organisms that contained complete 16S rRNA sequences (see methods section).

Therefore, in order to construct a more comprehensive sequence database we performed the taxonomic classification at the contigs-level. For this, we developed Python codes that enable to use protein sequences obtained from GTDB with DIAMOND and QIIME, to perform protein sequence alignment and classification of amplicon sequence variants, respectively (see methods section for Github repository of Python codes). A consensus lineage for each contig was determined using a modified version of the contig annotation tool (CAT) [79]. The stringency of the original CAT algorithm may result in a lower number of genus-level annotations. The algorithm and the parameters were therefore adjusted to improve genus level annotations, while adhering to taxonomies originally observed in the 16S amplicon sequencing experiments (SI-doc chapter 1, Figures S1–2). Moreover, to better standardize the taxonomic classification between metaproteomics, metagenomics and 16S amplicon sequencing, we established a ‘homogenized’ version of GTDB (SI-doc chapter 2). Advantageously, GTDB can be also employed to classify the 16S rRNA amplicon sequencing data because it contains small subunit ribosomal RNA sequences (ssu rRNA). However, approximately 15% of the representative taxa in GTDB contain 16S sequences that are shorter than 1200 base pairs and approximately 30% completely lack corresponding 16S sequences (SI-doc, chapter 2, Figure S3– 5). GTDB was therefore homogenized by selecting only taxa that are also represented with (full length) 16S rRNA sequences. Albeit this decreased the number of organisms in the constructed database, the comparative classification with the non-homogenized database showed only an overall decrease of approx. 5% of reads/PSMs matches (SI-doc Figures S6–8). Nevertheless, the homogenized database ensured a comparable reference sequence content and a therefore more accurate comparison between the approaches. Noteworthy, because the 16S-based classification follows a different principle it may therefore never be fully comparable to the contig-based classification (*e.g.*, Bayesian classifiers versus read assembly and sequence alignment).

### The impact of reference sequence database content divergences on obtained taxonomic profiles

The taxonomic classification of the metaproteomics and DNA and rRNA-based outputs was performed with a range of different reference sequence databases. The evaluation included protein sequence databases derived from UniProt (UniprotKB, UniRef, Swiss-Prot), NCBI (RefSeq) and ribosomal RNA reference sequence databases MIDAS and SILVA. Furthermore, we constructed a homogenized version of the recently-established genome taxonomy database (GTDB), which could be uniformly employed across the different approaches. The individual taxonomic profiles for the metaproteomics and the metagenomic data were visualized by Sankey diagrams to i) visualize the degree of annotation, the ii) flow in sequence annotations between databases, and to iii) evaluate the data for discrepancies within (or between) the different approaches (Figure 2). The graphs demonstrate highly-comparable phylum level profiles obtained after employing different reference sequence databases. At the lower taxonomic ranks, however, the application of different reference sequence databases led to substantial discrepancies in the obtained taxonomic profiles. For example, the UniprotKB and NCBI-derived RefSeq sequences that use the NCBI taxonomy showed a large number of (primarily) prokaryotic lineages that contain generic placeholder names and gaps characterized by keywords such as: ‘uncultured’, ‘unidentified’, ‘organism’, ‘metagenome’, ‘unknown’, ‘subgroup’, ‘group’, ‘bacterium’ or ‘proteobacterium’ (these are further referred to as ‘dump taxa’ in this study). For example, *Ca.* Accumulibacter is a prominent and key phosphate-accumulating organism in wastewater microbiomes. In GTDB, the lineage is: k__Bacteria, p__Proteobacteria, c__Gammaproteobacteria, o__Burkholderiales, f__Rhodocyclaceae, g__Accumulibacter. However, in NCBI taxonomy, *Ca.* Accumulibacter is stranded at the order and family level because of ‘uncertain placement’ where it only shows the gap annotation: ‘Betaproteobacteria incertae sedis’. Such inconsistencies strongly bias the taxonomic profiles and emphasizes the importance of rank normalization used in the genome taxonomy database. Moreover, as expected, increasing levels of sequence clustering reduces taxonomic resolution. Therefore, the highest degree of clustering resulted in a significant proportion of unnamed sequences. Likewise, the manually-annotated SwissProt database from UniProt Knowledgebase has a limited taxonomic coverage (approx. 300 K bacterial protein sequences, release 2021_03 statistics, https://web.expasy.org/docs/relnotes/relstat.html) and showed the lowest number of sequence annotations. Most importantly, for every metaproteomic experiment, a fraction of sequences remained that did not obtain any taxonomic classification, and which fraction could not be further interpreted. For GTDB, the fraction of sequences without annotations ranged from <5% at the phylum level, to approximately 25% at the genus level (this fraction is often also not shown in result graphs). In more extreme cases, *e.g.,* the highly-clustered UniRef sequence database, the fraction without genus-level annotation contributed to >50% of the total sequences. Overall, and as was expected, the Sankey graphs for the metaproteomic and metagenomic data followed very similar trends. The metagenomic data sets, however, showed a significantly greater diversity. This was also apparent in the ‘top 10 genera bar graphs’. For metagenomics, this fraction accounted for only around 5% of the total volume, compared to more than 33% for metaproteomics.

**Figure 2:**
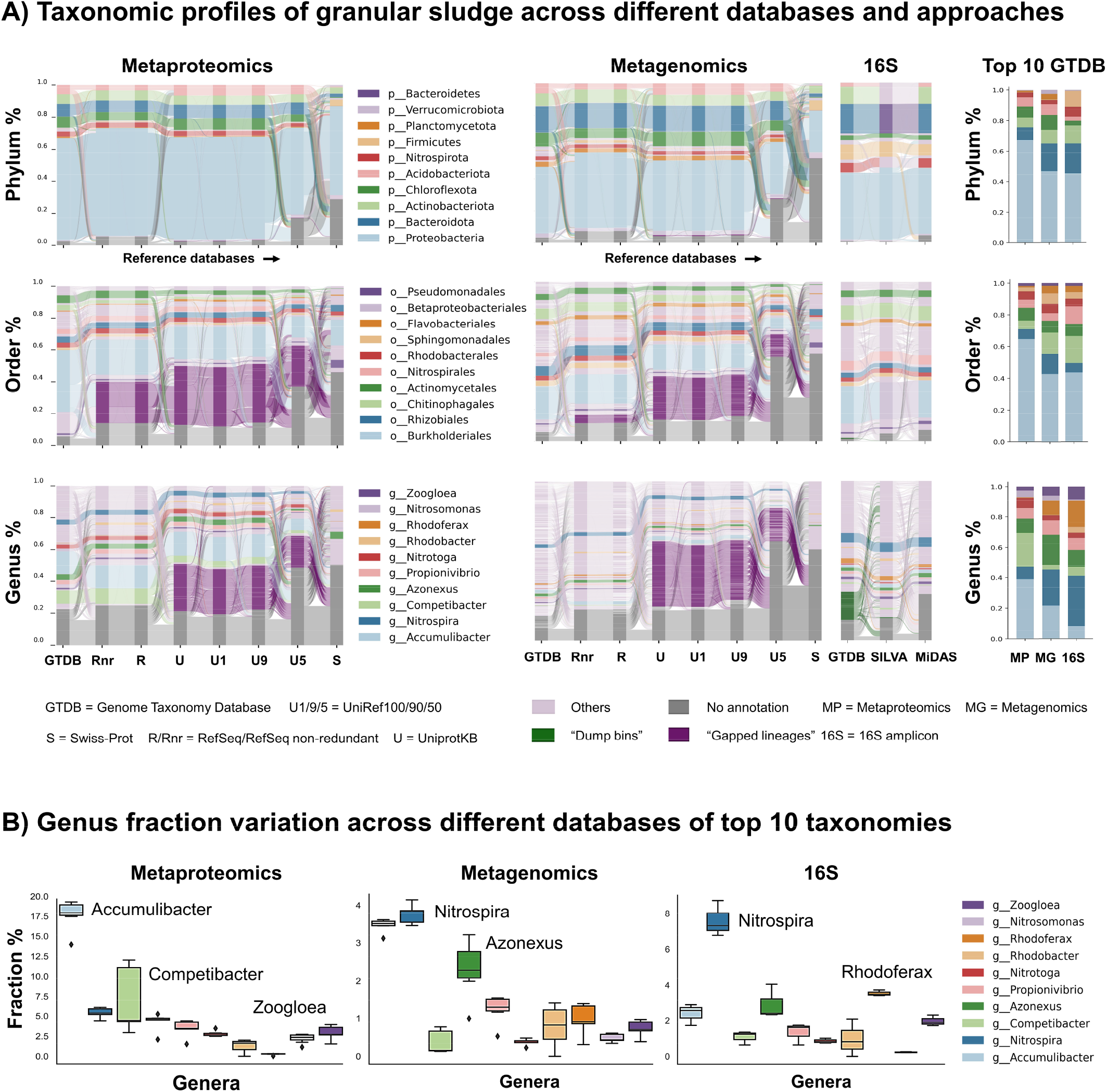
**A)** The Sankey flow diagrams show the impact of different reference sequence databases on the obtained taxonomic profiles for the granular sludge microbiome when using i) metaproteomics, ii) metagenomics, and iii) 16S amplicon sequencing (form the left to right). The applied reference sequence databases for classifying the metaproteomics and metagenomics data were GTDB, RefSeq non-redundant, RefSeq, UniprotKB, UniRef100, UniRef90, UniRef50 and SwissProt. The 16S amplicon sequencing data were classified using the ribosomal RNA reference sequence databases GTDB ssu reps, MiDAS and SILVA. The 10 most abundant taxa identified by the different approaches are furthermore presented as separate bar graphs on the right. Regardless the overall differences, the top taxonomies provide comparable profiles between the different approaches. GTDB represents a homogenized version of the Genome Taxonomy Database (GTDB) that contains only taxonomies with full length 16S reps. Taxonomic names were ‘unified’ as described in the methods section. Sankey flow diagrams that demonstrate the minor impact of the database homogenization on the overall microbiome coverage are shown in Figures S6–8. The Sankey flow diagrams and bar graphs were constructed by combining the annotations obtained from all aerobic granular sludge microbiomes (from plants 1–3). Extended Sankey flow diagrams for metaproteomics detailing all main taxonomic ranks are shown in Figure S9. **B)** The box plots show the genus fraction variation of the top 10 taxonomies for metaproteomics, metagenomics and 16S amplicon sequencing (from left to right) across the different reference sequence databases (excluding Swiss-Prot). Competibacter showed a considerably large variation in metaproteomics, where *Ca.* Competibacter, *Azonexus, Rhodobacer* and *Rhodoferax* on the other hand showed large genus fraction variations in metagenomics. The taxonomic abundances shown in the figures was determined by summing the total number of peptide-to-spectrum matches for metaproteomics, the ‘summed average depths of sequencing’ for metagenomics, and by using the total ASV counts for 16S rRNA amplicon sequencing.

Additionally, the GTDB-annotated 16S amplicon sequencing data were compared to the annotations obtained from SILVA NR99 and the wastewater-specific MiDAS databases (which both are based on the SILVA taxonomy framework). Thereby, a notable discrepancies were observed, e.g. for the genus *Tetrasphaera,* which is a key phosphate-accumulating organism (PAO) in wastewater treatment plants [19, 93]. When using the MIDAS and SILVA reference databases this genus appeared to be very abundant. Tetrasphaera was also observed when using the UniRef reference sequence database (albeit very low abundant), but it was nearly absent when using GTDB as reference sequence database. Interestingly, the same sequences were however found annotated with ‘c_Actinomycetia’. A closer inspection of GTDB sequences by BLAST + confirmed that although *Tetrasphaera japonica* was matched to each of the respective ASVs, it did not obtain the highest percentage identity for the 16S sequencing data. The best match however was a genus of the *Dermatophilaceae* family (SI-doc, Figure S10, and SI-EXCEL-4). This observation appears to be a limitation of the V3–V4 primer resolution and from a difference in phylogenetic placement because GTDB reassigns many taxa that are annotated as *Tetrasphaera* in NCBI to different genera. Complete tables with abundances as obtained for the individual databases can be found in the supplementary information (SI-EXCEL-5-7). Interactive Krona charts for the aerobic granular sludge microbiomes from the different wastewater treatment plants (1–3) across all approaches (classified by GTDB) are available via GitHub page https://pabstm.github.io/Comparative_metaproteomics_kronas/ and the supplementary information as excel macro-enabled workbooks (SI-EXCEL-8-16).

### Shared microbiome and biomass fractions between metaproteomics, metagenomics and 16S amplicon sequencing

When comparing the taxonomic profiles obtained by metaproteomics with those obtained by genomic approaches we observed differences already at the phylum level (Figure 2A). Here, the dominant proteobacteria encompass a larger fraction in metaproteomic experiments compared to the genomic methods. These differences are even more apparent at the lower genus level (Figure 2A and 2B). For example, *Ca.* Competibacter, *Ca.* Accumulibacter and *Ca.* Nitrotoga are very prominent in the metaproteomic experiments; but are much less pronounced in the metagenomic and 16S amplicon sequencing data. On the other hand, *Nitrospira* appears (relatively) abundant when performing genomic experiments, and the 16S amplicon data moreover suggest a high abundance of *Tetrasphaera* and *Rhodospherax.*

Nevertheless, regardless of the differences in the overall taxonomic annotations, the relative proportions of the top 10 taxonomies between the different approaches (when using the ‘homogenized’ GTDB) were surprisingly comparable (Figure 2A, right bar graphs). Nonetheless, the total number of taxonomies (genera) that were observed by all three techniques appeared relatively moderate (Figure 3A, Venn diagrams: 52, 39 and 47 genera for plants 1–3, respectively). On the other hand, the ‘total abundance fraction’ which those shared genera covered was comparatively large (Figure 3A, grey bars, lower bar graphs). For example, the genera that were observed by all three approaches accounted in metaproteomics for approximately 80% of the total abundance. The abundance fraction the shared genera covered in 16S amplicon sequencing was however significantly lower (approx. 30–60%, depending on the treatment plant). Because metagenomics and 16S amplicon sequencing showed a large number of very low abundant taxonomies, for this evaluation only genera that were observed by at least two techniques and that were >3% of the total abundance were considered. Taxonomies that express central nutrient-removing genes were included regardless of their abundance.

**Figure 3:**
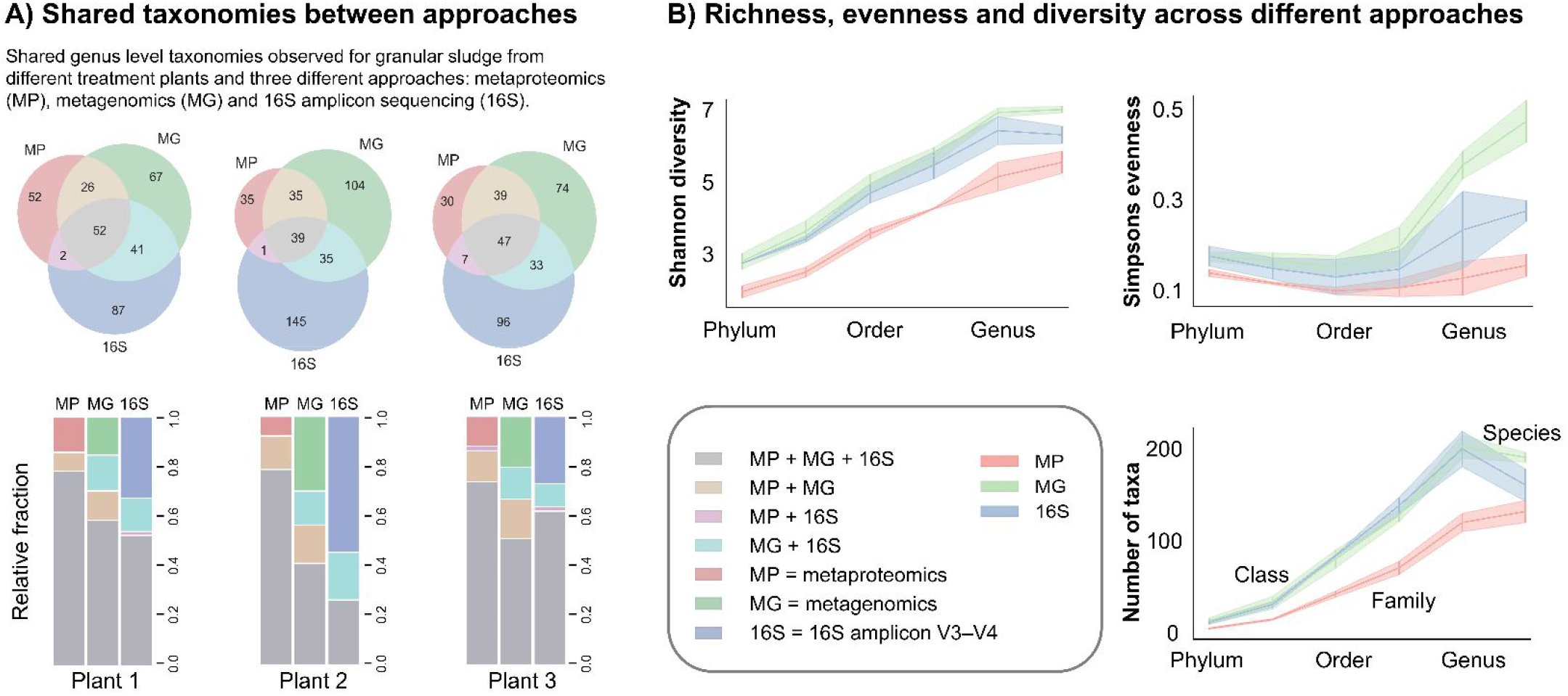
**A)** Shared genus-level taxonomies between metaproteomics and the DNA and rRNA-based approaches for the granular sludge microbiomes from the three wastewater treatment plants 1–3. Data is expressed as numbers of shared taxa (upper images, Venn diagrams) or as a fraction of total shared abundance fraction (lower images, bar graphs). The graphs consider only taxonomies that were observed by at least two techniques (non-unique taxa), and which taxa further were present at >3% abundance (compared to total abundance of non-unique taxa), or which express central nutrient-removing genes. Albeit the fraction of genera that were uniformly observed by all 3 approaches appears relatively moderate (grey sections, Venn diagrams), those taxonomies coverage the majority of the protein-biomass in metaproteomics (grey bars, bar graphs, labelled with ‘MP’). The color codes are further described in the box. **B)** The graphs visualize the microbial diversity indices i) ‘Richness’, ii) ‘Simpson’s Evenness’ and iii) ‘Shannon diversity’ for the granular sludge microbiomes as obtained from the different approaches. The graphs are displayed for the taxonomic ranks phylum, class, order, family, genus, and species. The data from the three plants (1–3) were averaged. Graphs in A and B were generated by using a homogenized Genome Taxonomy Database (GTDB) that contains only taxonomies with ‘full length’ 16S reps. As expected, the DNA and rRNA based approaches (green and blue traces) show a significantly larger number of genera, but suggest also a much higher taxonomic evenness, compared to metaproteomics (red traces).

Furthermore, both metagenomics and 16S amplicon sequencing generally showed a larger diversity, richness and evenness compared to metaproteomics (Figure 3B). 16S amplicon sequencing, for example, identified the largest number of taxonomies at the genus level (approx. 200). This was not unexpected, as the DNA and rRNA-based approaches utilize amplification steps and also measure free genetic material, dead and dormant microbial cells. On the other hand, albeit metaproteomics appears to have a lower sensitivity and therefore identified the lowest number of genera, the shared taxonomies (considering the above mentioned thresholds) accounted for a large fraction of the measured (protein-based) biomass.

### Observed key metabolic genes and metabolic traits

Two key processes of nutrient removal in wastewater treatment are the elimination of nitrogen and phosphorous. To assess genera involved in the conversion of these two core processes we integrated the functional annotations obtained from KEGG, COG terms, PFAM, TIGRFAM domains and UniprotKB genes (Figure 4). Interestingly, the functional gens covering the nitrogen processes are currently fragmented across different databases. For example, *Nxr* annotation was annotated via UniprotKB, *Nap* by KEGG, and *Nar* using COG terms. Polyphosphate-accumulating organisms (PAO) remove phosphate from the wastewater by producing polyphosphate with the genes *ppk* (polyphosphate kinase) and *ppa* (pyrophosphatase). Glycogen-accumulating organisms (GAO) – that compete with PAOs for short-chain fatty acids – synthesize glycogen using *glg* (glycogenin glucosyltransferase) and likely therefore show also high expression of *bglX* (beta-glucosidase like enzymes). Nitrogen removal is achieved via subsequent nitrification and denitrification steps that is performed by *hao* (hydroxylamine oxidoreductase) and *amo* (ammonia monooxygenase) genes of ammonia-oxidizing bacteria (AOB) and *nxr* (nitrite oxidoreductase) of nitrate-oxidizing bacteria (NOB). Denitrification (DN) is encoded by the gene clusters *nar* (respiratory nitrate reductase) and *nap* (periplasmic nitrate reductase) to reduce nitrate and *nirK* (copper-containing nitrite reductase) and nirS (cytochrome cd1-containing nitrite reductase) to reduce nitrite, while the genes *nor* (nitric oxide reductase), *nrf* (nitrite reductase) turnover nitric oxide, and ultimately, *nos* (nitric oxide synthase) converts nitrous oxide to dinitrogen gas. Cyc (cytochrome C) is implicated in either the activity of *nor* or *nrf.* Interestingly, *nor* proteins were only detected at low levels, which supposedly is a consequence of membrane association or of poor database annotation accuracy. Furthermore, *hzs* (hydrazine synthase), *hdh* (hydrazine dehydrogenase) as well as *hao* (hydroxylamine oxidoreductase) could be detected in one plant, which are part of the anammox process such as found in *Ca.* Brocadia. Interestingly, several of the key nutrient-removing genera appeared very low abundant in metagenomics and 16S amplicon sequencing data, which was in contrast to the metaproteomics outcomes. These include genera such as *Accumulibacter, Competibacter* and *Propionivibrio* (PAO, GAO and DN, respectively), *Nitrosomonas* (AOB) and *Nitrotoga* (NOB and DN), and *Zoogloea* (DN). Conversely, several other genera, such as *Azonexus* (PAO and DN) and *Nitrospira* (NOB and DN) showed only a minor difference between the orthogonal methods. In addition to *Sulfuritalea* (PAO and DN), other genera were even more prominent in the DNA and rRNA-based approaches. For *Ca.* Accumulibacter, this observation is in agreement with previous studies [57, 94, 95], but for *Ca*. Competibacter, however, the observed differences have not been reported before. Moreover, a recent large-scale genomic study showed the widespread presence of genes such as *nosZ* (nitrous-oxide reductase) or *ppk* (polyphosphate kinase), which were detected in a large fraction of the MAGs [96]. However, *ppk* for example, could be actually observed by metaproteomics in only a few genera at significant levels. The search terms (used in this study) to extract functional information from the metaproteomics data, as well as a complete table detailing protein taxonomic and functional annotations for all treatment plants can be found in the supplementary information (SI-EXCEL-17-18).

**Figure 4:**
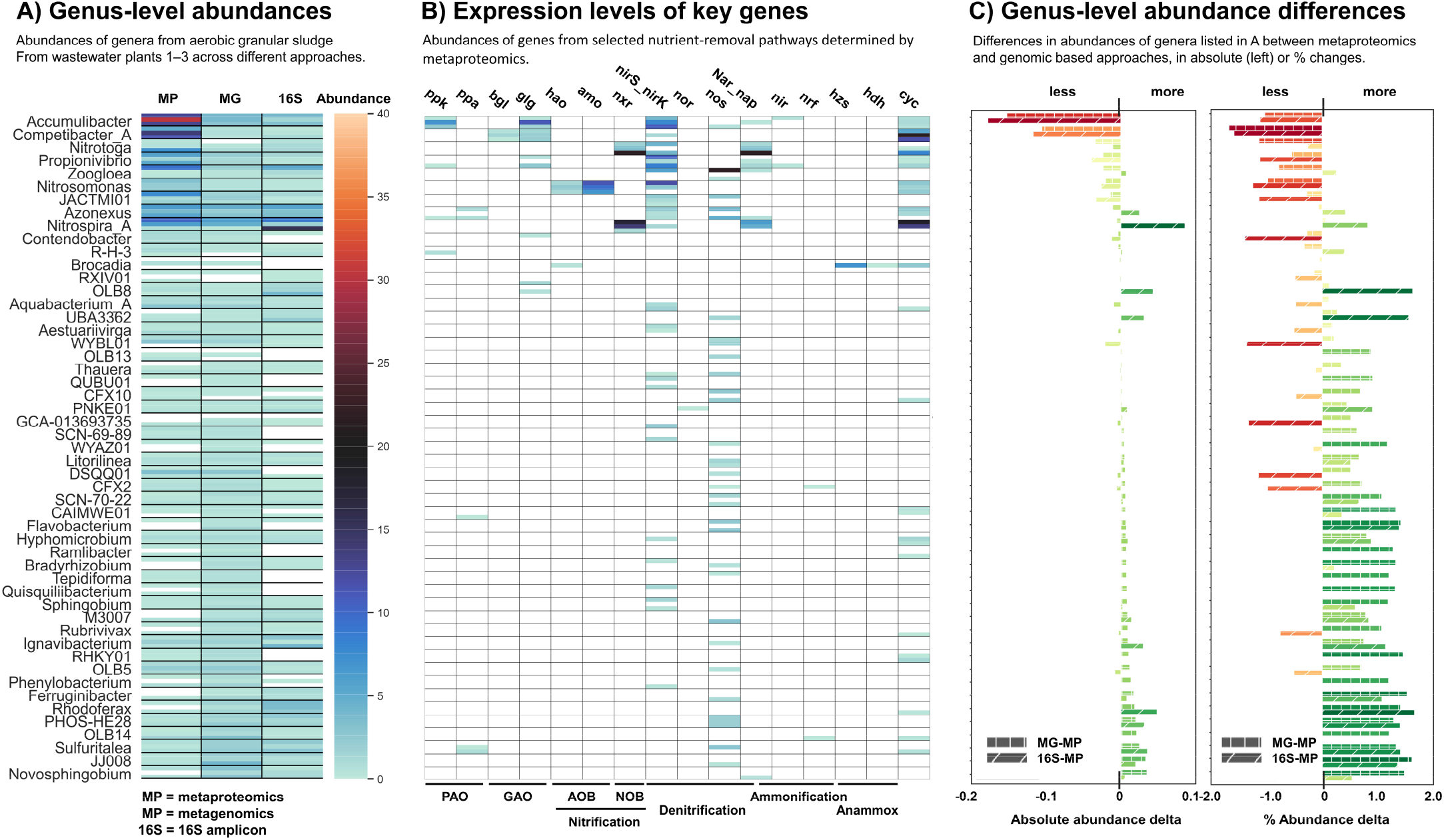
**A)** The heat map shows (key) genera that are present in the aerobic granular sludge microbiome, and which are potentially involved in the central nutrient-removal processes that take place during the wastewater treatment. The taxonomic abundances observed in metaproteomics (MP), metagenomics (MG) and 16S rRNA amplicon sequencing (16S) are shown in separate columns (from left to the right). The abundances observed in the microbiomes obtained from the different treatment plants are shown as individual bars within one cell (top bar = plant 1, middle bar = plant 2 and lower bar = plant 3). Generally, the most dominant genera observed in metaproteomics are *Ca.* Accumulibacter followed by *Ca.* Competibacter. In metagenomics, the most abundant genera are *Nitrospira, Ca.* Accumulibacter and *Azonexus.* Nevertheless, for the genomic approaches, taxonomies were generally found more evenly distributed. **B)** The heat map details expression levels of genes from selected nutrient-removal pathways as observed by metaproteomics. The genes are named on the top of the heat map (ppk = polyphosphate kinase, ppa = pyrophosphatase, bglX = beta-glucosidase-like, glg = glycogenin glucosyltransferase, hao = hydroxylamine oxidoreductase, amo = ammonia monooxygenase, nxr = nitrite oxidoreductase, nirK = copper-containing nitrite reductase, nirS = cytochrome cd1-containing nitrite reductase, nor = nitric oxide reductase, nos = nitric oxide synthase, nar = respiratory nitrate reductase, nap = periplasmic nitrate reductase, nir = nitrite reductase genes (converting nitrite to nitric oxide), nrf = nitrite reductase (which converts nitrite to ammonium), hzs = hydrazine synthase, hdh = hydrazine dehydrogenase and cyc = cytochrome). The corresponding pathways or organisms are indicated below the heat map (PAO = phosphate accumulating organism, GAO = glycogen accumulating organism, AOB = ammonia-oxidizing bacteria, NOB = nitrite oxidizing bacteria) **C)** The graph shows the abundance differences for the selected genera between metaproteomics and metagenomics (bars with vertical pattern), or metaproteomics and 16S amplicon sequencing (bars with diagonal pattern). The differences are expressed as absolute differences (left plot) or as % abundance differences (right plot). Graphs were generated by using a homogenized Genome Taxonomy Database (GTDB) that contains only taxonomies that are also represented by ‘full length’ 16S reps.

### Categorization of the observed metaproteome

The COG classification system was used to further categorize the observed proteins and to group them into the categories ‘metabolism and transport’, ‘membrane-associated’, ‘cell cycle’, and ‘other’ (Figure 5, and a detailed table with annotated proteins can be found in SI-EXCEL-18). Approximately 80% of the peptide-to-spectrum matches could be grouped into any of the categories following the combined annotation using KEGG, PFAM and TIGRFAM. Comparing the obtained peptide-to-spectrum matches with the contig-based frequencies provided furthermore a measure of enrichment between metaproteomics and metagenomics. Next to nutrient removal (carbohydrate, nitrogen, amino acid metabolism) and growth (translation) also membrane-associated proteins were highly-prominent. An interesting observation was the strong enrichment of porin proteins. These are a beta barrel-forming class of transporters expressed in gram-negative organisms and included fatty acid transporters (*fadL*), small inorganic molecules (*cirA, fepA, ovp1*) and coenzyme transports (*btuB*). The presence of enriched amounts of membrane proteins, however, did not directly correlate to a potential bias in abundance. For example, the genera *Ca.* Competibacter and *Ca.* Propionivibrio showed strong differences in abundance between metaproteomics and metagenomics but showed little difference in profiles derived from peptide-to-spectrum matches and contig-based frequencies. On the other hand, genera such as Aquabacterium_A and *Azonexus* showed highly-enriched fractions of membrane proteins, but only small differences in relative abundance between the protein and the DNA-based approach. *Ca.* Accumulibacter had an increased difference with increasing porin protein content (highest in wastewater plant 2), while *Ca.* Competibacter from wastewater plant 1 contained a smaller porin fraction and showed a smaller difference. Species-level differences between samples showed predominantly *Ca.* Accumulibacter phosphatis_G from wastewater plant 2, *Ca.* Accumulibacter phosphatis_C from wastewater plant 1 and 3, and a reduced fraction of *Ca.* Competibacter denitrificans from wastewater plant 1. Therefore, observed differences are presumably rather related to phenotypes and species-level variations. For 16S rRNA amplicon data, biases are well-researched and attributed to variations in 16S rRNA gene copy number and primer choice. For example, the increased gene copy number presumably resulted in an overestimation of Firmicutes, while a reduced copy number may have underestimated the abundance of Acidobacteriota and Verrucomicrobiota. The V3–V4 primers used in this study did not efficiently amplify *Chloroflexeota* which consequently were only observed at low levels. Planctomycetota were not captured by 16S rRNA amplicon sequencing at all.

**Figure 5:**
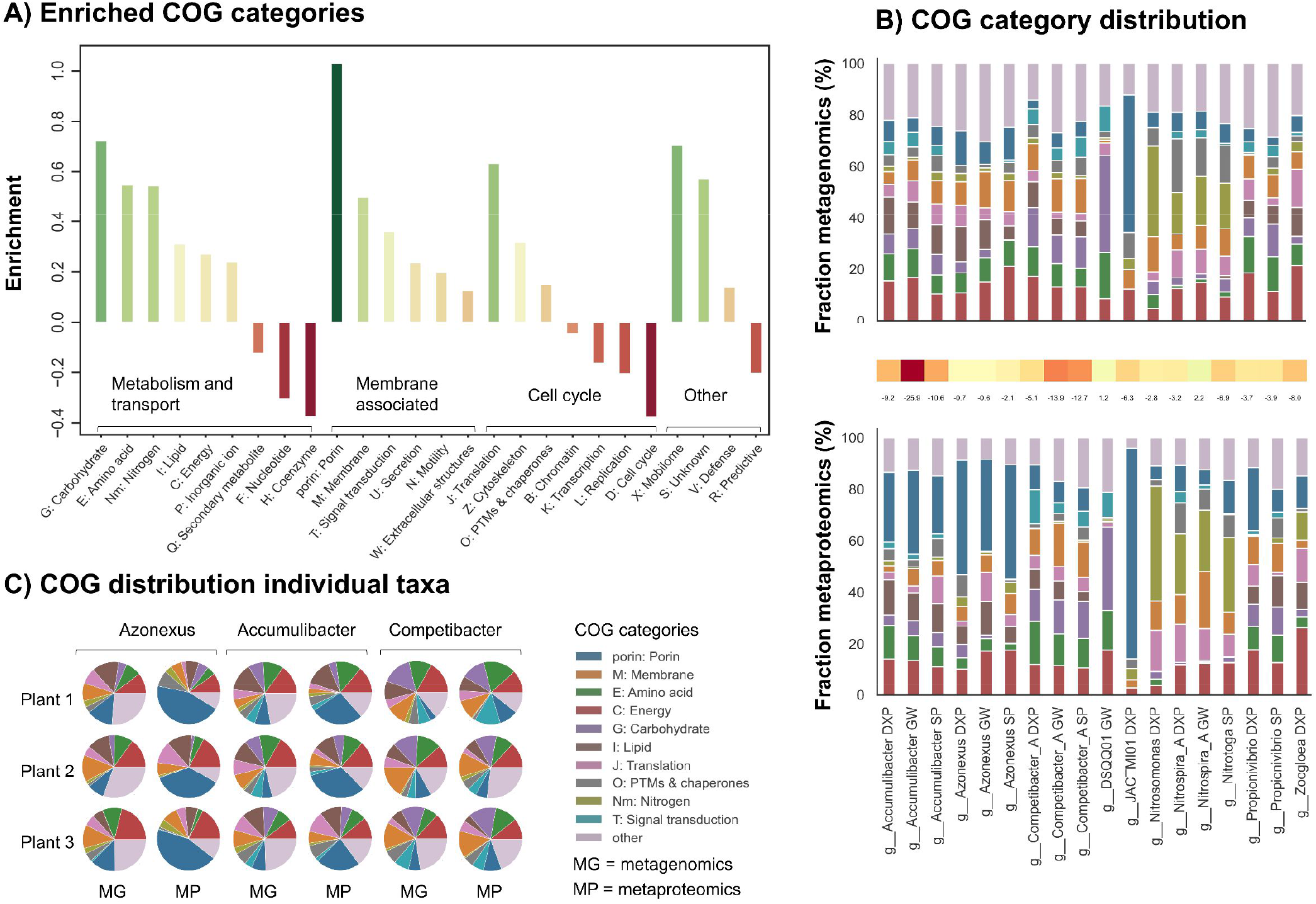
**A)** The bar graph shows COG term enrichment analysis (Clusters of Orthologous Groups) for the metaproteomics data of the combined (averaged) granular sludge microbiome data. B) The graph compares COG category distribution of abundant organisms between metagenomics (upper graph) and metaproteomics (lower graph). C) Comparison of proportion of COG categories for selected organisms between metaproteomics (MP) and metagenomics (MG). Metagenomics data are based on the summed depth of sequencing while metaproteomics data are based on peptide-to-spectrum matches (PSMs). The graphs show averaged data from the wastewater plants 1–3 (except otherwise stated).

## Discussion and conclusions

The application of large-scale ‘omics’ approaches, such as metaproteomics and metagenomics, are increasingly used in microbial ecology and biotechnology. Therefore, efforts have been devoted to standardizing methods in the fields of metagenomics, 16S amplicon sequencing, and metaproteomics. For example, these efforts resulted in the CAMI study for metagenomics [59] and more recently to the CAMPI study for metaproteomics [58]. Both studies aimed to compare methodologies and outcomes between laboratories. While microbiome studies that integrate different types of approaches are rapidly evolving, studies that systematically investigate the orthogonal character of metaproteomics and genomic approaches have rarely been performed [35, 45, 97]. Among the sources that significantly impact variability between studies and approaches are the different reference sequence databases and the employed taxonomies that are used by the different approaches. Database content divergences and inconsistencies, as well as inaccurate taxonomies and nomenclatures can profoundly impact the accuracy of the taxonomic representation and comparisons between studies and techniques. Here, we report on a comparative metaproteomic characterization of the aerobic granular sludge microbiome. We show the divergent views on the central nutrient-removal organisms that can be obtained depending on the chosen ‘omics’ approach and reference sequence databases. Additionally, we demonstrate the uniform application of a ‘homogenized’ genome taxonomy database, which enabled a more accurate interpretation of the orthogonal nature of metaproteomics and the genomic approaches. Ultimately, the performed study demonstrates the importance of metaproteomics for the characterization of complex microbiomes and the application of accurate and uniform reference sequence databases to enhance comparative studies and scientific reporting.

Python codes for constructing and formatting Genome Taxonomy Database (GTDB) entries for the use with Diamond and QIIME are openly accessible via https://github.com/hbckleikamp/GTDB2DIAMOND and https://github.com/hbckleikamp/GTDB2QIIME. Interactive Krona charts showing the aerobic granular sludge microbiomes from the different wastewater treatment plants as obtained for the different omics approaches are available via the GitHub page https://pabstm.github.io/Comparative_metaproteomics_kronas/.

## Supporting information

SI EXCEL OVERVIEW

SI DOC

## ACKNOWLEDGEMENTS

The authors acknowledge Carol de Ram for support with sample preparation, Claudia Tugui and all other colleagues from the department of Biotechnology for valuable discussions, Leanne van Benthem for collection of the granule materials, and the SIAM (Soehngen Institute of Anaerobic Microbiology) consortium for funding.

## CONFLICT OF INTEREST

The authors declare no competing interests.

## ABBREVIATIONS

MG: metagenomics (whole metagenome sequencing)
MP: metaproteomics
16S: 16S rRNA gene sequencing
AGS: aerobic granular sludge
DXP: Dinxperlo, plant 1 (wastewater treatment plant, The Netherlands)
GW: Garmerwolde, plant 2 (wastewater treatment plant, The Netherlands)
SP: Simpelveld, plant 3 (wastewater treatment plant, The Netherlands)
ASV: amplicon sequence variant
PSM: peptide-to-spectrum match
MAG: metagenome assembled genome
GTDB: genome taxonomy database
PAO: phosphate-accumulating organisms
GAO: glycogen-accumulating organisms
AOB: ammonium-oxidizing bacteria
AOA: ammonium-oxidizing archaea
NOB: nitrate-oxidizing bacteria
NR: nitrate-reducing organisms
EPS: extracellular polymeric substances

## Notes

### Competing Interest Statement

The authors have declared no competing interest.

