## Supplementary material for "Comparative metaproteomics demonstrates different views on the complex granular sludge microbiome": SI DOC

### SI information material to:

**Table of contents**

1. Adjusted CAT (contig annotation tool) approach
2. Database (GTDB) homogenization procedure
3. Impact of GTDB homogenization procedure on taxonomic profiles
4. Sankey graphs for metaproteomics for all databases and taxonomic rankings
5. Sequence homology alignment for Tetrasphaera sequences
6. Additional references

### 1. Adjusted CAT (contig annotation tool) approach

The taxonomic classification of contigs was performed according to the CAT approach ('contig annotation tool', which was originally published by Meijenfeldt et al., 2019 [1]. This approach firstly infers the taxonomic affiliation of ORFs from the lowest common ancestor (LCA) of the best hit lineages from DIAMOND sequence alignments ('protein LCA'). In the following a consensus lineage for each contig is constructed from the annotated ORFs by a 'contig-level LCA'. CAT was developed to provide improve classification precision in particular for sequences from unknown organisms, thus the (stringency of the) approach results in a rather conservative number of genus-level annotations. Therefore, we adjusted the CAT approach with the objective to increase the taxonomic resolution, which appears justified because the wastewater microbiome is in the reference sequence databases comparatively well covered. The major modification of the original approach includes a 'stepwise' construction of a lineage starting from superkingdom to species level. Thereby, the best scoring taxon of a rank is determined only from the group of taxa that is part of the 'best scoring taxonomic lineage' of the higher ranks (Figure S1). The total bitscore (and consequently also the minimal bit score support which defines the cutoff to assign a taxon to a contig) is recalculated on every taxonomic rank. This increases the number of genus-level assignments without impacting annotations of higher taxonomic ranks.

Furthermore, the adjusted approach includes filtering for contigs based on the size. The original approach does not suggest any considerations based on the contig length. Short contigs, however, can more easily introduce false positive classifications, in particular if only a few ORFs are (or a single ORF is) covered by the contig. The adjusted approach, therefore, includes a filtering step to filter taxa that were only annotated to contigs with less than a given number of predicted ORFs (e.g.<5).

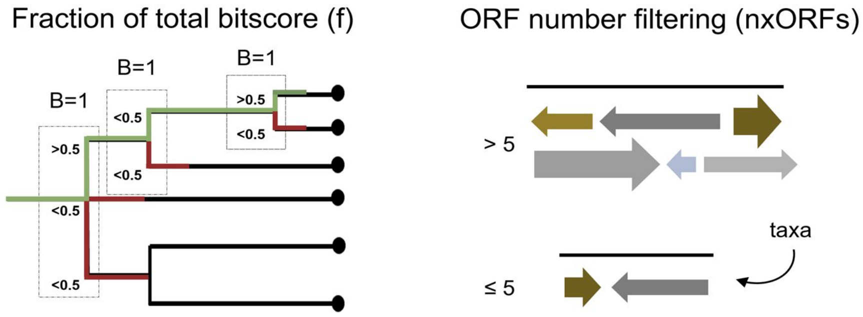


**Figure S1.** The graphs shows the introduced adjustments to the original CAT approach published by Meijenfeldt et al., 2019 [2]. The adjusted approach includes a 'stepwise' construction of a lineage, where the best scoring taxon per rank is determined only from the group of taxonomies that are part of the 'best scoring taxonomic lineage' of the higher ranks. Therefore, the minimal bit score support (which is the cutoff to assign a taxon to a contig) is recalculated at every taxonomic rank (left graph). After the taxonomic annotation, a filtering step was included to filter taxa that were only found in contigs with less than a given number of predicted ORFs (e.g.<5).

Finally, a parameter sweep was performed to determine the best combination of ‘f’ (which corresponds to the fraction of the sum of bitscores, and that determines the minimal bit-score support ‘mbs’) and ‘nxORF’ (which corresponds to the minimum required number of ORFs per contig), which is shown in Figures S2. The parameter sweep was performed with the objective to maximize the Jaccard similarity coefficient to the 16S amplicon sequencing data (see materials and methods section). As baseline comparison, the original CAT algorithm parameters r=10 and f=0.5 were used [1], in which ‘r’ denotes the range of top bitscore % used to determine the lowest common ancestor (LCA) of ORFs, and ‘f’ to the fraction of the sum of bitscores, as described above. In addition we included two different approaches to determine the lowest common ancestor for the individual ORFs (protein LCA), which were the standard approach (LCA, r=10) and a ‘top bitscore’ approach (BLCA, r=0).
The final selected parameters used for taxonomic classifications within the current study are ‘BLCA’ for protein LCA with the adjusted contig LCA (fraction of bitscore LCA = FbLCA, f=0.5, nxORFs=5) for the classification of contigs. Using the optimized approach and parameters, the genus level annotations improved from 0.27 to 0.82; while the Jaccard similarity coefficient (to the 16S amplicon data) improved from 0.19 to 0.25 (Figure S2).

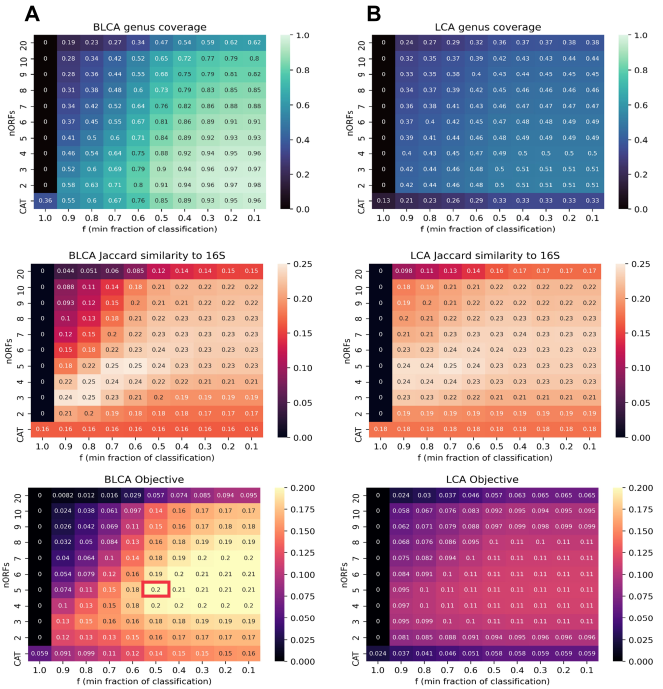


**Figure S2.** The heat maps show the performed parameter sweep for: i) the fraction of bitscore (f) and ii) the minimum number of ORFS (nxORFS) using the adjusted contig-LCA (FbLCA) combined with two different protein LCA options ‘BLCA’ (graphs A) and ‘LCA’ (graphs B). A parameter sweep for ‘f’ (0.1–1) and ‘nxORF’ (2–10 and 20) was used to find the best combination of parameters. As an objective score, a multiplication was used of the genus coverage and the Jaccard similarity coefficient compared to taxonomies observed in the 16S amplicon data. The best performing combination of the BLCA with the contig LCA parameters f=0.5 and nxORFs=5 (or 4) is highlighted by a red square (left lower graph).

### 2. Database (GTDB) homogenization procedure

16S rRNA genes are challenging to include into assembled genomes because of the diverging GC-content and varying copy numbers and the discrepancies between databases. Additionally, different databases also contain different length distributions. For example, Silva ssu Parc is less curated compared to the Silva NR99, where many short, fragmented sequences are found. MiDAS, however, only contains full-length sequences [3].

GTDB contains next to genome and protein sequences also small subunit 16S ribosomal RNA databases: i) reps, which is 'representative', and ii) all, which contains all sequences [4]. However, when comparing GTDB protein reps to 16S reps (Figure S3), approx. 30% of genomes in GTDB protein reps do not have a corresponding 16S sequence (labelled with 'MG only'), and approx. 15% of sequences are shorter than 1200 base pairs, which was selected as the cut-off for full-length sequences in this study. Comparing GTDB protein reps to "16S all", showed several genomes that were not present in GTDB protein reps (labelled with '16S only') that consequently would also lead to database discrepancies. Approx. 75% of all phyla have one representative in the 16S rRNA database (Table S1). Figure S4 moreover shows the fraction of representation of each individual phylum, where several candidate phyla with provisionary names do not have a single 16S rRNA representative. On the other hand, common phyla such as 'Proteobacteria' are much stronger represented. Figure S5 further demonstrates how database discrepancies are distributed over the taxonomic tree, where for example 'Bacteroidetes' and 'Firmicutes_A' are phyla that typically contain higher 16S rRNA gene copy numbers.

**Table S1.** Coverage indicates which percentage of taxa for each taxonomic rank contains at least one representative in the 16S reps database. Sequences shorter than 1200 base pairs were considered as fragmented, while longer sequences were considered as full length sequences.

| target | full length | fragmented | MG only | coverage (%) |
|---:|---:|---:|---:|---:|
| genomes | 26031 | 6853 | 15010 | 54.35 |
| superkingdom | 2 | 0 | 0 | 100.0 |
| phylum | 121 | 7 | 24 | 79.61 |
| class | 290 | 29 | 71 | 74.36 |
| order | 721 | 139 | 239 | 65.61 |
| family | 1578 | 474 | 761 | 56.1 |
| genus | 5185 | 1741 | 2893 | 52.81 |
| species | 17753 | 4202 | 8618 | 58.07 |

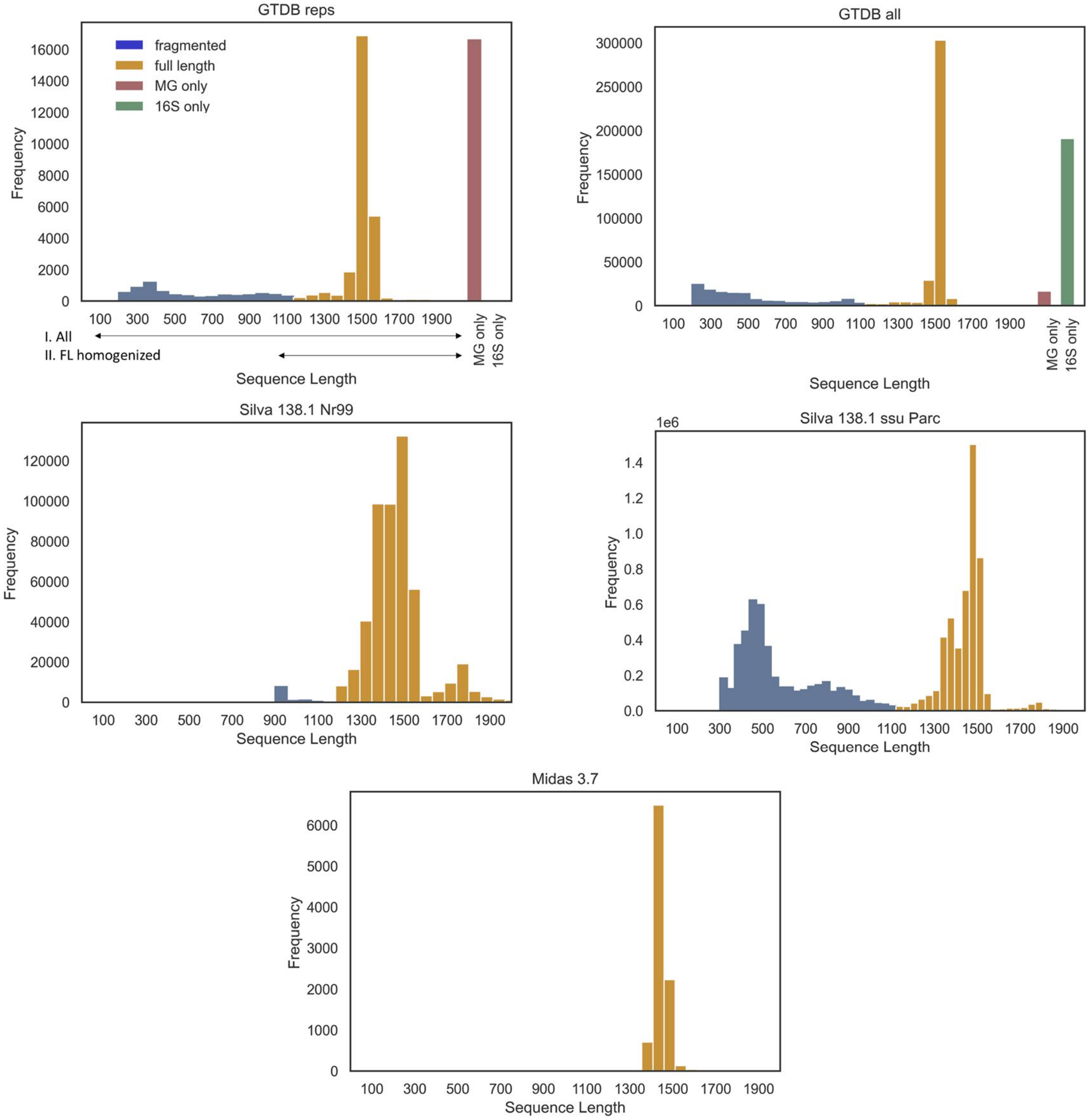


**Figure S3.** The graphs show the 16S rRNA length distributions determined for different reference sequence databases. The taxonomies found for GTDB '16S reps' and '16S all' was compared to the GTDB protein reps database. Sequences that were only present in GTDB protein reps were labelled with 'MG only' and entries that were only present in the 16S database were termed '16S only'.

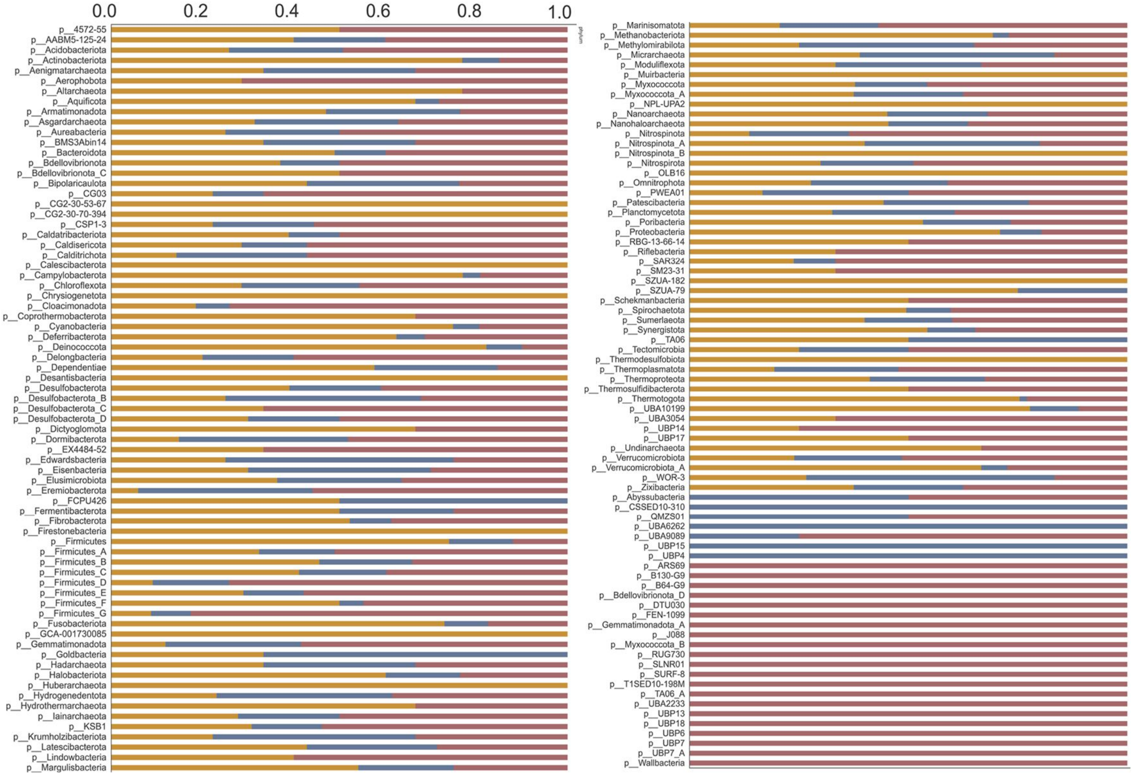


**Figure S4.** Coverage of phyla in the GTDB protein reps database compared to GTDB 16S reps sequences. Coverage is based on the percentage (1 is equal to 100%) of species within the phylum that are represented in GTDB 16S reps. The fraction of fragmented 16S sequences that are shorter than 1200 base pairs are shown as blue bars, while > 1200 base pairs are treated as full length sequences and are shown as yellow bars. A complete lack of representation are shown as red bars.

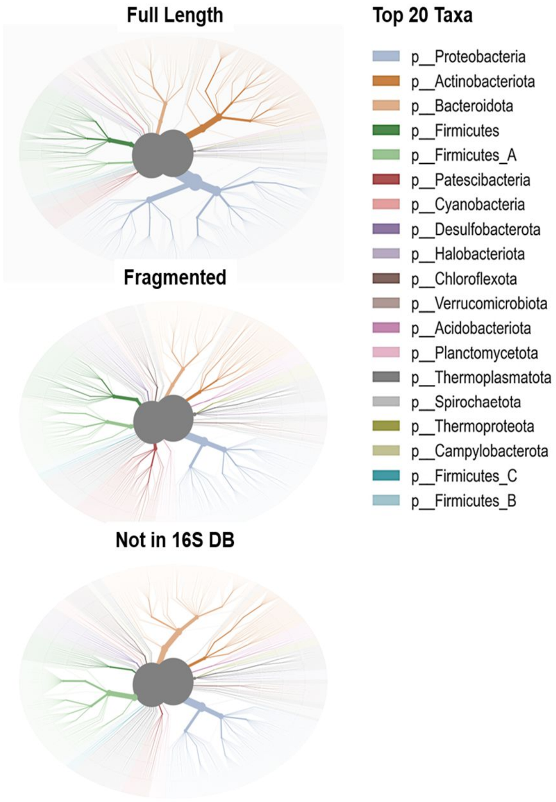


**Figure S4.** Representation of taxa over taxonomic ranks (nodes are in the order of domain, phylum, class, order, family, genus to species) within the taxonomic tree of the GTDB protein reps genomes, that either contain a full length 16S representative (>1200 base pairs), a fragmented representative (<1200 base pairs) or have no representative (not in 16S). The top 20 taxonomies are indicated by different colors.

### 3. Impact of GTDB homogenization on taxonomic profiles

The effect of the GTDB homogenization procedure (as described in chapter 2) was furthermore evaluated using Sankey diagrams. Overall, the homogenization showed limited impact on the observed taxonomic profiles (~5% of genus level changed for the metaproteomics profiles) and were only more pronounced at the species level.

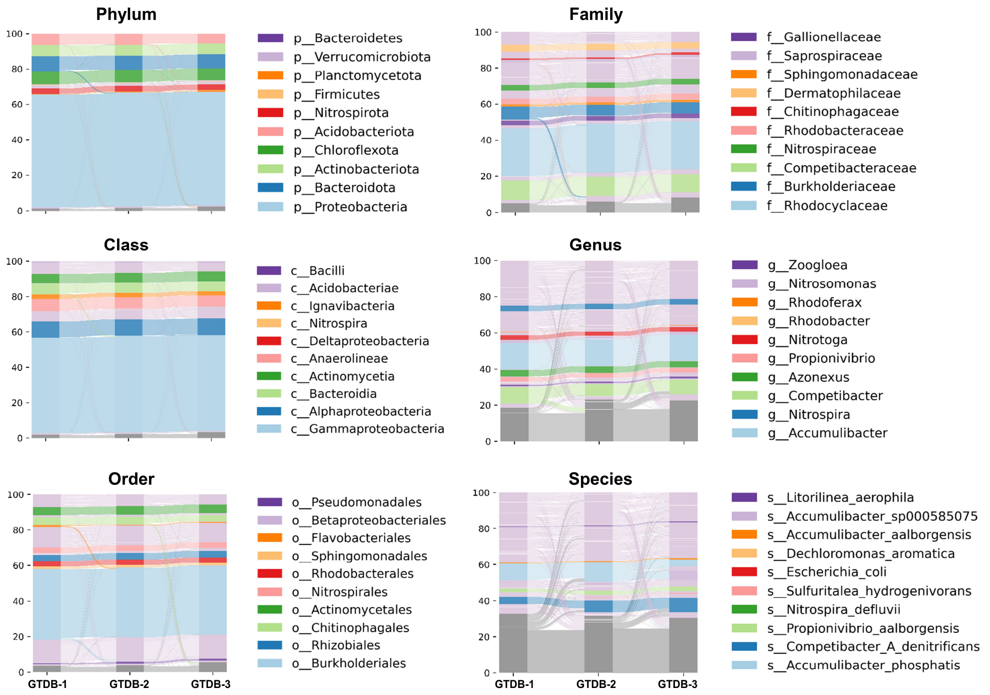


**Figure S6.** Metaproteomic taxonomic profiles displayed as Sankey diagrams of the combined microbiomes from plants 1–3, over the ranks phylum to species. Taxonomic classification was performed with GTDB protein sequence that were homogenized at different levels. GTDB-1: GTDB reps (all) non-homogenized, GTDB-2: protein reps sequences homogenized to contain those present in 16S representative ssu sequences and GTDB-3: GTDB protein reps sequences homogenized to contain those present in 16S representative ssu sequences larger than 1200 bp.

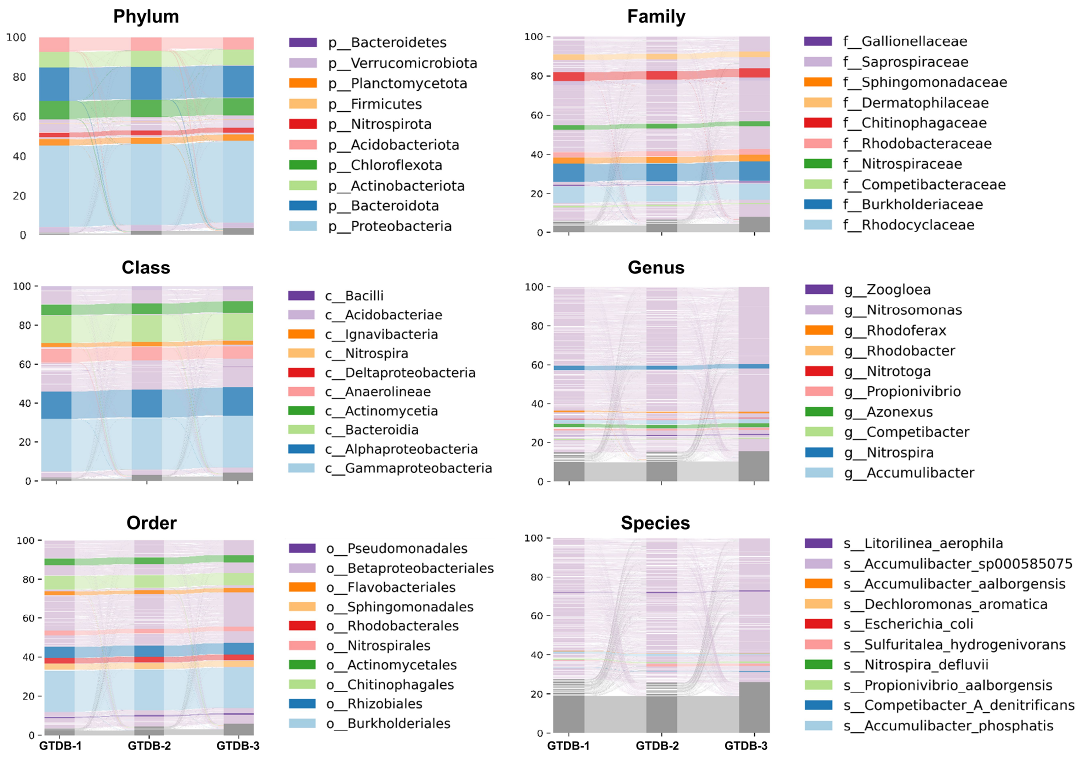


**Figure S7.** Metagenomic taxonomic profiles displayed as Sankey diagrams of the combined microbiomes from plants 1–3, over the ranks phylum to species. Taxonomic classification was performed with GTDB protein sequence that were homogenized at different levels. GTDB-1: GTDB reps (all) non-homogenized, GTDB-2: protein reps sequences homogenized to contain those present in 16S representative ssu sequences and GTDB-3: GTDB protein reps sequences homogenized to contain those present in 16S representative ssu sequences larger than 1200 bp.

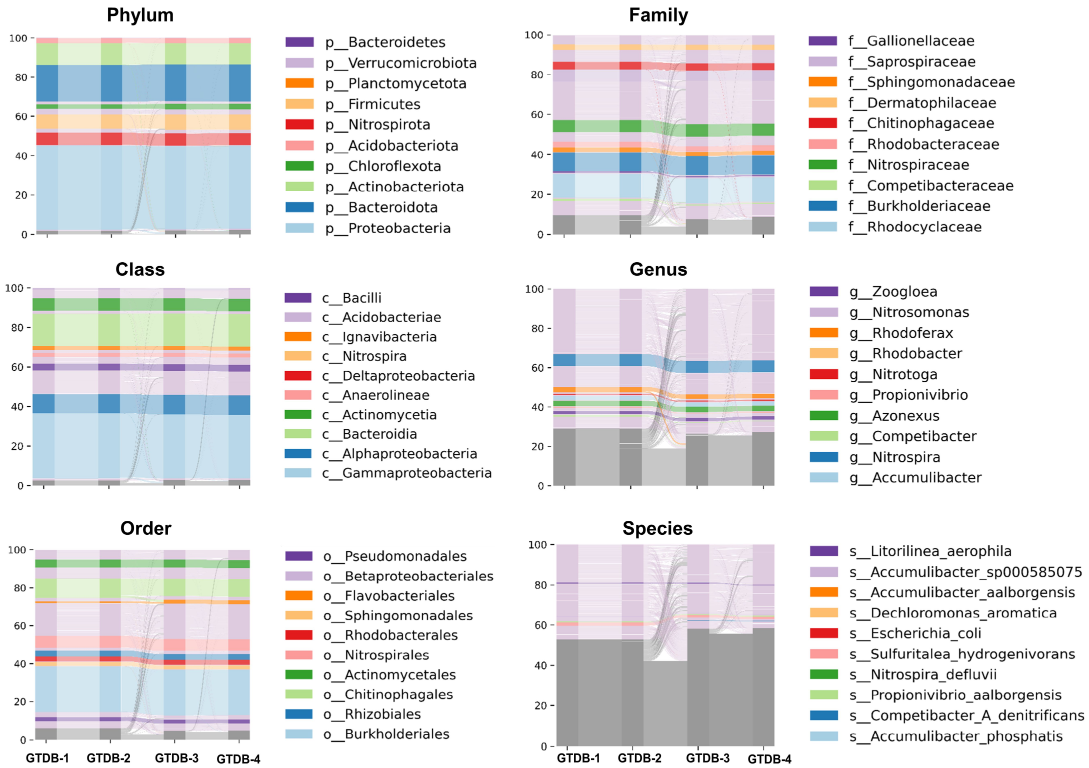


**Figure S8.** 16S amplicon sequencing-based taxonomic profiles displayed as Sankey diagrams of the combined microbiomes from plants 1–3, over the taxonomic ranks phylum to species. Taxonomic classification was performed with GTDB SSU rRNA sequences that were homogenized at different levels. GTDB-1: GTDB reps (non-homogenized), GTDB-2: GTDB representative sequences longer than 1200 bp, GTDB-3: GTDB all sequences (non-homogenized) and GTDB-4: GTDB all sequences longer than 1200 bp.

### 4. Sankey graphs for metaproteomics for all databases and taxonomic rankings

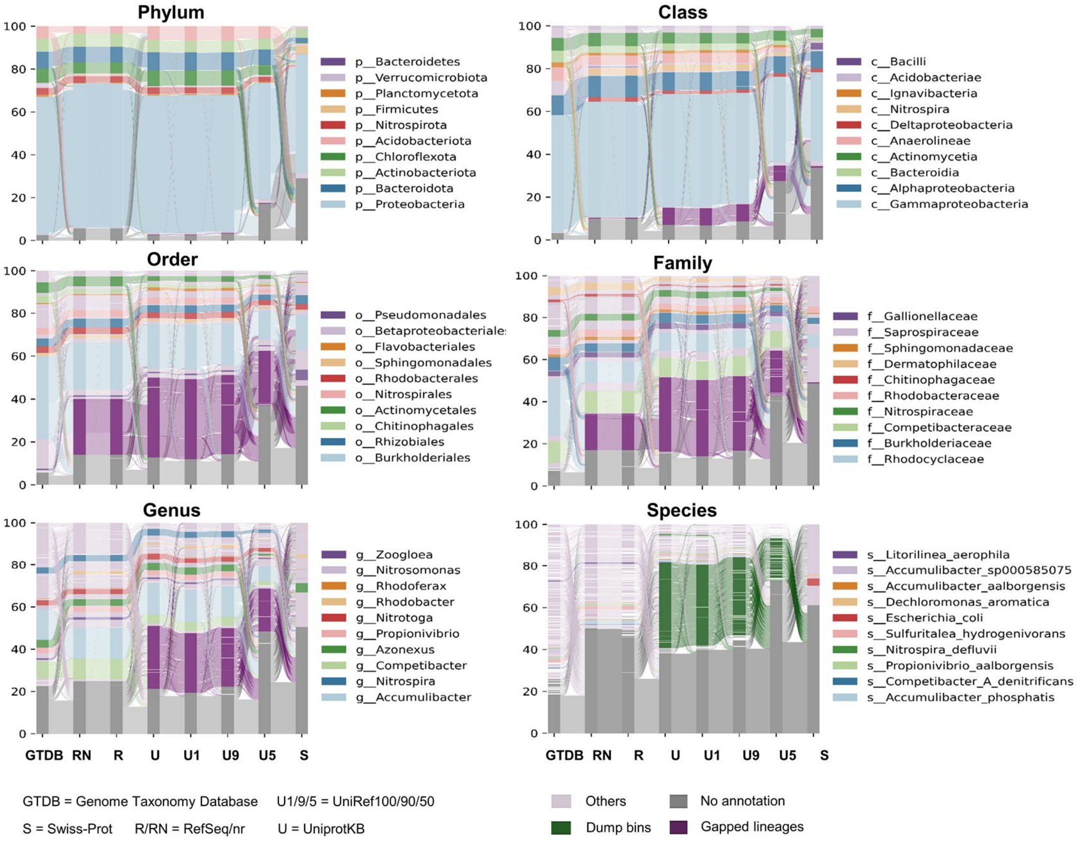


**Figure S9.** Sankey flow diagrams demonstrating the impact of different reference sequence databases on the observed taxonomic profile for the granular sludge microbiome (averaged over plants 1–3) analysed by metaproteomics. The individual graphs detail the main taxonomic ranks Phylum – Species. The y-axis represents the abundance of the respective taxonomy as fraction of the total (100%). The top taxonomies are named in the legends and are shown in individual colors. The profiles obtained from applying the different reference databases is plotted along the x-axis. The employed reference sequence databases are (from left to right): GTDB, RefSeq non-redunant, RefSeq, UniprotKB, UniRef100, UniRef90, UniRef50 and Swiss-Prot. GTDB represents a homogenized version of the Genome Taxonomy Database (GTDB), which contains only taxonomies with full length 16S reps.

### 5. Sequence homology alignment for Tetrasphaera sequences

The absence of Tetrasphaera was a surprising difference between GTDB ssu, Silva and Midas annotations. Therefore, ASVs were aligned using Blastn to GTDB FL (full length, >1200 base pairs) ssu. For Silva, 3 ASVs were annotated with Tetrasphaera (Figure S14A) and for Midas, 20 (Figure S14B). The alignment revealed that within GTDB, several genera show better identity percentages to other genera (not annotated with Tetrasphaera). Therefore, when using GTDB, Tetrasphaera appears less represented.

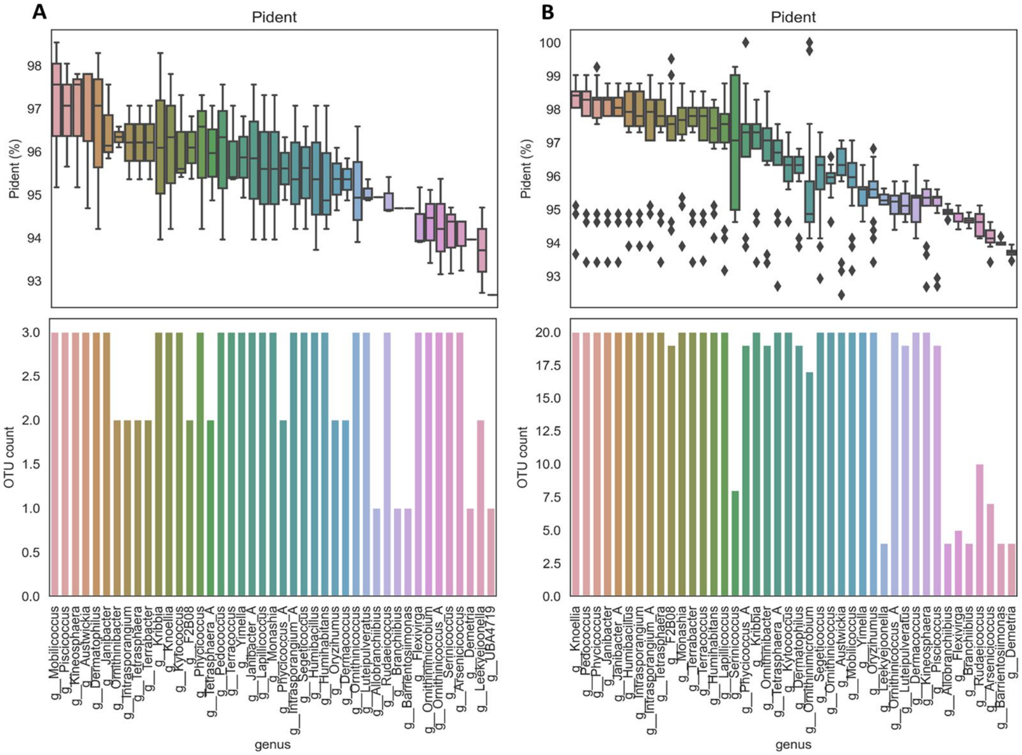


**Figure S10.** Tetrasphaera ssu annotations for GTDB. A) distribution of GTDB genera aligned with Blastn for Silva ASVs with family Dermatophilaceae, B) distribution of GTDB genera aligned with Blastn for Midas ASVs with family Intrasporangiaceae.
